# Characteristics of human donor lungs utilized for research

**DOI:** 10.1101/2024.08.10.607431

**Authors:** Nishanth R Shankar, Lorena Garnica, Emily Stahlschmidt, Derek Byers, Hrishikesh Kulkarni

**Affiliations:** Kasturba Medical College, Mangalore, India 575001; Division of Pulmonary and Critical Care Medicine, Washington University School of Medicine, St. Louis, MO 63110; Mid-America Transplant, St. Louis, MO 63110

## Abstract

**Introduction:** Many fundamental discoveries have occurred using primary cells from deceased donor lungs. These cells respond differently to injury when there are underlying co-morbidities like diabetes mellitus, hypertension, aging and exposures to cigarette smoke, cocaine and chronic alcohol use. However, the prevalence of these characteristics in donor lungs utilized for research is currently unknown.

**Methods:** This retrospective cohort study procured data of lung transplant donors from Mid-America Transplant from January 2017 until July 2023. The donors were characterized based on lung utilization into three groups - lungs used for research, lungs used for transplant, and lungs not recovered from donors for either research or transplantation.

**Results:** The mean age of donors whose lungs were utilized for research was 41±18 years. 25% of them were expanded criteria donors (ECD) while 10% of the donors in the transplant cohort were ECD. 14% of the donors whose lungs were utilized for research had history of diabetes compared to 8% of donors whose lungs were transplanted. A quarter of the research donor population had positive history of cigarette use within the preceding 20 years. At least 40% of donors had a positive history of non-intravenous drug use, of whom a majority had a history of continued non-intravenous drug use.

**Conclusions:** No strict selection criteria or protocols exist when human donor lungs are obtained for ex-vivo research. There is a high prevalence of diabetes mellitus, history of smoking and non-intravenous drug use along with older age distribution in donors whose lungs used are for research.

## INTRODUCTION

Tissues from organ donors that are not used for transplantation provide a valuable resource for investigating human disease and are used as complementary models to animal studies. Many fundamental discoveries have been conducted using primary cells derived from deceased donor lung tissue. [(1),(2)] These primary cells behave differently to injury and inflammation when there are underlying co-morbidities like diabetes mellitus (DM), hypertension (HTN), aging and exposures to cigarette smoke, cocaine and chronic alcohol use. [(3),(4)] For example, cultured differentiated bronchial epithelial cells derived from brushings obtained from people with alcohol use disorder had altered barrier function and proinflammatory responses compared to those without alcohol use disorder. [(5)] Lungs from these donors with diabetes have differential responses to infection. [(6)] However, the prevalence of these characteristics in donor lungs that are not utilized for transplantation but are utilized for research, is currently unknown. In this current study, we aimed to study the prevalence of various co-morbidities and exposures in lung donors. We also aimed to determine the prevalence of certain characteristics like age distribution of donors, exposure to cigarette smoke and relevant co-morbidities like DM in donors whose lungs were used for research is necessary because the outcome of various ex vivo studies conducted on deceased human lung tissue may be dependent on them. Moreover, knowledge gained from this work will form the basis for power calculations required for research studies that utilize donor lungs, for example, those investigating pneumonia. Based on our observations, we conclude that accounting for various co-morbidities in donors of lungs utilized for research is important in designing ex vivo studies and interpreting their results.

## METHODS

### Study design

We conducted a retrospective cohort study of transplant donors based on their lung disposition.

### Study setting

We procured data of all lung transplant donors amounting to a total of 1686 donors with various outcomes at Mid-America Transplant from January 2017 until July 2023. Mid-America Transplant is a non-profit organization based in St. Louis, Missouri which works with its partner hospitals spread across 84 counties in Missouri, Illinois, and Arkansas to procure organs and provide them to recipients in need across their designated service area and beyond. The lungs were used for research by academic institutions who had an agreement with Mid-America Transplant (including Washington University). The lungs that were transplanted were utilized by Mid-America Transplant partner hospitals. Since the data procured was of deceased human donors, this study did not qualify as human subjects research under 45CFR46.102(e)(1).

### Primary outcome of lung transplant donors

The lungs were characterized based on their predisposition into three groups - lungs used for research, lungs used for transplant, and lungs which were not recovered from donors. The data procured contained extensive patient information including demographics (age, race, gender, blood group), anthropometry (weight, height, BMI), cause and manner of death (mechanism of death, circumstance of death, organ donor type), medical history (history of diabetes mellitus, hypertension, coronary artery disease, previous myocardial infarction, cancer) drug exposure and social history (history of intravenous (IV) drug use, high risk behavior, cigarette use in the past 20 years, cocaine use, non-IV drug use), serologies (Hepatitis B, Hepatitis C, Human Immunodeficiency Virus, Cytomegalovirus), recent chest radiograph readings and recent microbiological culture data.

### Donor variables

Donors who had documented history of diabetes mellitus (DM) prior to the current hospitalization for donation were included as positive for DM. Those who were prescribed insulin for management of diabetes were included under the criteria of “Insulin dependent diabetes mellitus”. Similarly, patients who had documented history of hypertension (HTN) were included under “Positive for HTN”. Patients with documented diagnoses of coronary artery disease (CAD) or myocardial infarction (MI) in their medical records were included to be “Positive for history of CAD” and “Positive for history of MI” respectively. History of cigarette usage equal to or exceeding 20 pack-years were included in “positive history for cigarette use of 20 pack-years” while history of cigarette use in the 6 months prior to their death in addition to history of smoking of more than 20 pack-years was considered as inclusion criteria for “cigarette use in the last 6 months”. A pack-year is defined as number of cigarette packs smoked per day multiplied by number of years smoked. [(7)] Extended criteria donors (ECD) were defined as brain dead donors who are over 60 years old, or donors who are 50-59 years old and have two or more of the following - history of high blood pressure, creatinine level of 1.5 mg/dl or higher, as measured by a blood test that shows kidney function, and/or death from a stroke or cardiovascular accident (CVA).

High-risk behavior was present if the deceased donor has factors associated with an increased risk for disease transmission, including blood-borne pathogens and if the deceased donor meets the criteria for increased risk for HIV, Hepatitis B, and Hepatitis C transmission set forth in the current U.S. Public Health Services (PHS) Guideline. Donors were included under “History of Cocaine use” if the donor has ever abused or had a dependency to cocaine and “History of continued cocaine use” was present if the donor abused or had a dependency to cocaine within the last 6 months. Donors were included under “history of non-IV drug use” if the donor had ever abused or had a dependency to Non-IV street drugs, such as crack, marijuana or prescription narcotics, sedatives, hypnotics or stimulants and “history of continued non-IV drug use” were people who abused or had a dependency to non-IV street drugs, such as crack, marijuana or prescription narcotics, sedatives, hypnotics or stimulants within the last 6 months. Donors included under “IV drug use” were donors who had ever abused or had a dependency to Intra-venous drugs which were not prescribed and used for non-medical reasons.

### Chest radiographs and scoring

Descriptive reports of the most recent chest radiographs were obtained for all the donors. For comparative analysis of chest radiographs between the three groups of donors, we adapted a scoring system which is a part of a lung donor scoring system used to predict outcomes in transplant recipients, which assigned clear radiographs as 0, minor changes as 1, opacity in ≤ 1 lobe as 2 and opacity in > 1 lobe as 3. [(8)] Individual chest radiograph description for every donor was analyzed and converted to a numeric code in accordance with the above-mentioned scoring system.

### Microbiological data

Most recent microbiological culture data of lung donors which included cultures of sputum, bronchoalveolar lavage/washings and blood was procured from Mid-America Transplant. A donor was characterized depending on the type of gram-positive bacteria, gram-negative bacteria, fungi, or virus being positive in culture. Additionally, we also categorized donors based on their positive results of blood cultures for bacteria and fungi.

### Statistical analysis

Data of the three donor categories (lungs utilized for research, transplant and lungs rejected) was analyzed independently for further comparison under common parameters such as demographics, anthropometry, medical history, drug exposure and personal history, cause, and manner of death, serologies, chest radiograph score and microbiological culture data using IBM SPSS Statistics Version 29.0.2.0.

## RESULTS

### Donor demographics and death characteristics

The mean age of donors utilized for research was found to be 41±18 years while that of donors whose lungs were utilized for transplant was found to be 35±14 years. Majority of the donors were of white/Caucasian decent under all three categories (Table 1).

**Table 1.** Donor demographics.

| **Characteristics** | **Research Lungs**<br>**(n = 128)** | **Transplanted Lungs**<br>**(n = 391)** | **Rejected Lungs**<br>**(n=1167)** |
| --- | --- | --- | --- |
| **Age** |  |  |  |
| Mean | 41.04±18 | 34.86±14 | 42.29±17 |
| Median | 41.5 | 33 | 44 |
| Interquartile range | 29 | 21 | 24 |
| **BMI\*** |  |  |  |
| Mean | 28.52±9 | 26.92±6 | 29.04±9 |
| Median | 26.76 | 25.86 | 27.38 |
| Interquartile range | 8.9 | 6.79 | 9.88 |
| **Race** |  |  |  |
| White/Caucasian | 71.88% | 74.17% | 77.72% |
| Black/African American | 24.22% | 21.99% | 19.36% |
| Hispanic | 2.34% | 2.55% | 1.29% |
| Others | 1.56% | 1.27% | 1.60% |
| **Gender** |  |  |  |
| Male | 56.25% | 59.59% | 62.98% |
| Female | 43.75% | 40.40% | 37.01% |
| **Blood Group** |  |  |  |
| O | 50% | 48.59% | 45.15% |
| A | 38.26% | 41.42% | 38.03% |
| B | 10.15% | 8.69% | 12.68% |
| AB | 1.56% | 1.28% | 4.11% |

\*Abbreviations: BMI – Body Mass Index.

The leading cause of death among all donor groups was found to be anoxia. However, the proportion of donors for research lungs who died from anoxia was double compared to those who died of head trauma or a stroke. In comparison, the proportion of donors for transplanted lungs who died from anoxia was nearly similar to those who died of head trauma (Table 2). Intracranial hemorrhage or stroke was the leading mechanism of death in donors used for research at 19% while drug/intoxication was the leading mechanism of death in donors used for transplant at 23%.

**Table 2.** Donor death characteristics.

| **Characteristics** | **Research Lungs**<br>**(n = 128)** | **Transplanted lungs**<br>**(n = 391)** | **Rejected lungs**<br>**(n=1167)** |
| --- | --- | --- | --- |
| **Cause of death** |  |  |  |
| Anoxia | 51.56% | 39.39% | 52.70% |
| Head Trauma | 22.66% | 38.87% | 22.62% |
| Cerebrovascular / Stroke | 20.31% | 20.72% | 20.65% |
| Other | 5.46% | 1.02% | 4.02% |
| **Death type** |  |  |  |
| Brain Death | 70.31% | 95.14% | 60.33% |
| Asystole | 29.69% | 4.86% | 39.67% |
| **Mechanism of death** |  |  |  |
| Intracranial hemorrhage/<br>Stroke | 18.75% | 20.72% | 20.91% |
| Cardiovascular | 17.19% | 5.63% | 19.11% |
| Blunt Injury | 17.19% | 22.51% | 17.99% |
| Asphyxiation | 11.72% | 5.12% | 7.71% |
| Drug / Intoxication | 10.16% | 23.27% | 18.68% |
| Gunshot wound | 7.03% | 17.39% | 6.08% |
| Others | 17.97% | 5.36% | 9.48% |
| **Circumstances of death** |  |  |  |
| Death from Natural Causes | 53.91% | 30.18% | 50.04% |
| Accident, Non-MVA* | 18.75% | 31.19% | 25.36% |
| Motor Vehicle Accident | 10.94% | 15.09% | 12.43% |
| Alleged Suicide | 10.16% | 16.11% | 6.86% |
| Alleged Homicide | 4.68% | 6.91% | 3.68% |
| Others | 1.56% | 0.51% | 1.63% |
| **Organ donor type** |  |  |  |
| SCD* | 46.88% | 85.17% | 41.05% |
| DCD* | 25.13% | 4.85% | 39.33% |
| ECD* | 25% | 9.72% | 19.45% |
| Null | 0% | 0.26% | 0.72% |

\*Abbreviations: Non-MVA – Non – Motor Vehicle Accidents; SCD – Standard Criteria Donor; DCD – Donor After Circulatory Death; ECD – Expanded Criteria Donors.

It is worthwhile to note that the percentage of standard-criteria donors (SCD) was only 47% in donors whose lungs were used for research as compared to 85% in donors whose lungs were transplanted (Table 2). Donation after cardiac death (DCD) donors were more prevalent among those whose lungs were used for research at 25% while only 5% of the donors whose lungs were transplanted were of DCD type. A higher prevalence of expanded criteria donors (ECD) was noted in those used for research with 25% of the cohort being ECD, while only 10% of the donors in the transplant cohort were ECD.

### History of non-communicable disease

14% of the donors whose lungs were utilized for research had history of DM, which was comparable to rejected lungs (Table 3). In comparison, only 8% of the donors whose lungs were transplanted had history of DM. High rates of prevalence of hypertension were found in donors used for research (41%), whereas only 23% of the donors whose lungs were transplanted were found to have history of hypertension (Table 3). The rate of coronary artery disease (CAD) in donors whose lungs were used for research was nearly 2.5 times of donors whose lungs were transplanted (Table 3).

**Table 3.**
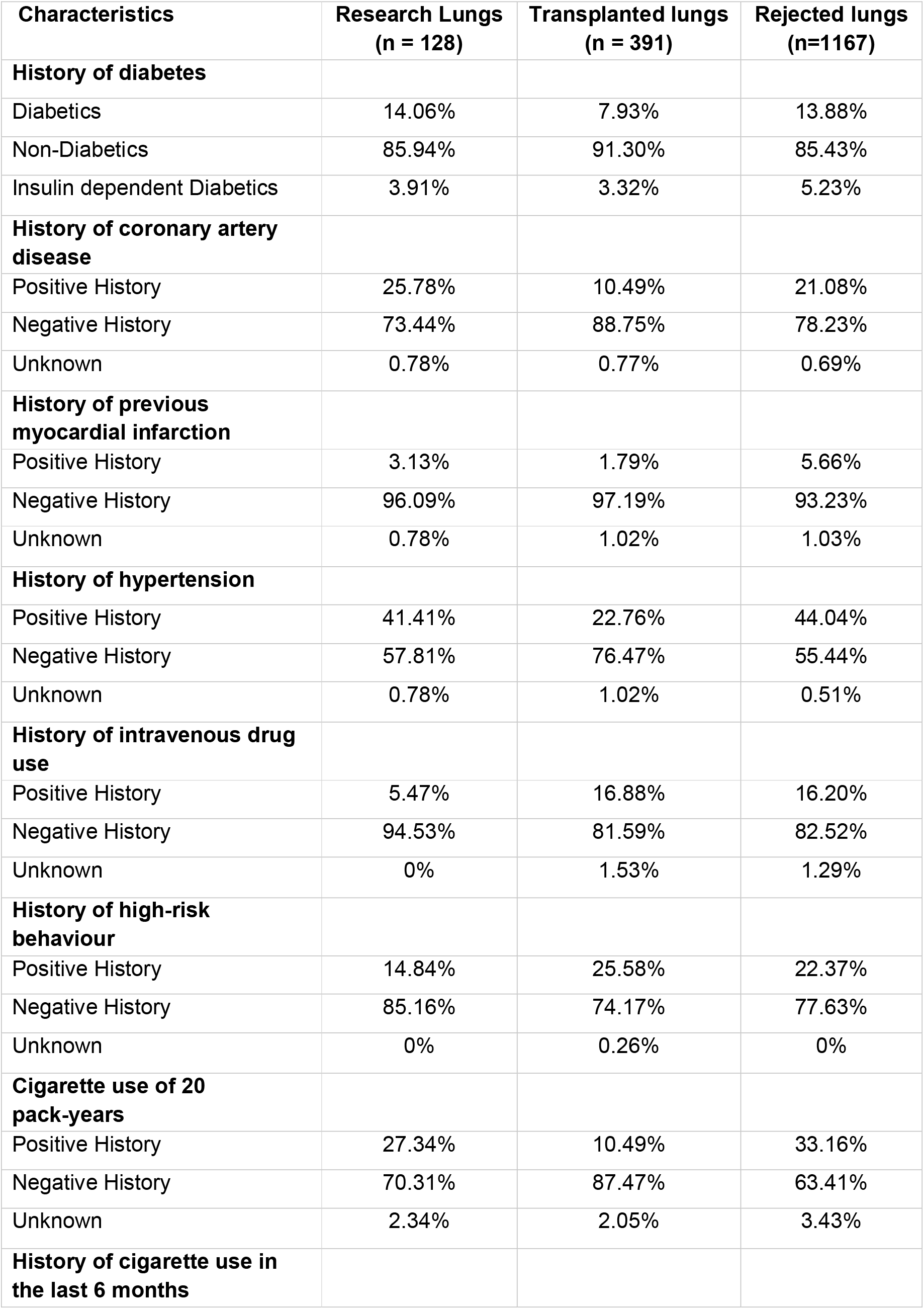
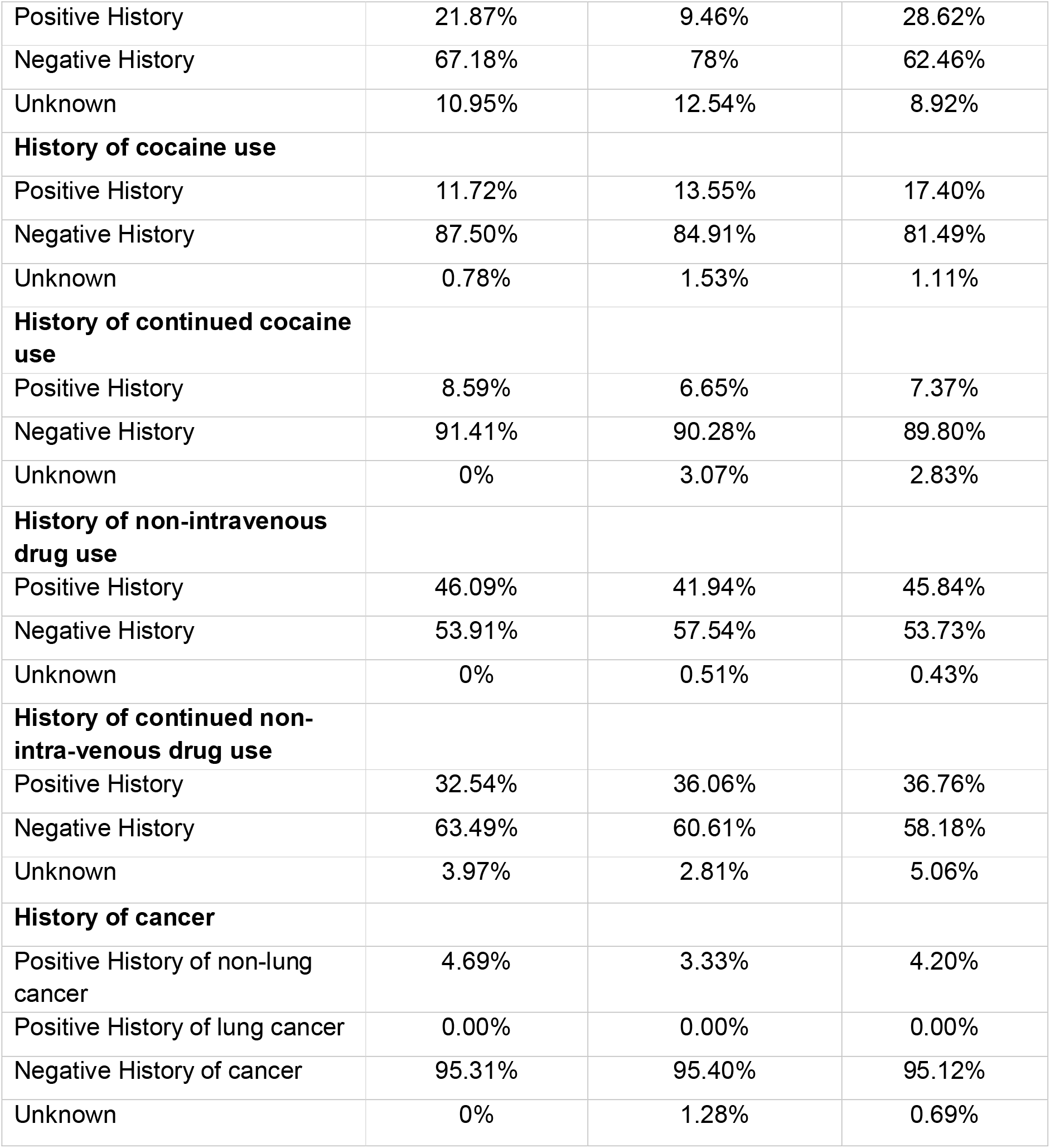
Donor history of non-communicable diseases, exposures, and cancer.

### Exposure history

A quarter of the research donor population had positive history of cigarette use exceeding 20 pack-years, whereas it was noted in only 10% of the donors whose lungs were transplanted (Table 3). In comparison, only 5% of the donors whose lungs were used for research had history of IV drug use, compared to 17% of the donors used for transplant (Table 3). History of cocaine use was similar between the two groups. Of note, at least 40% of the donors were found to have a positive history of non-IV drug use, of whom a majority had a history of continued non-intravenous drug use (Table 3).

### Serology

<1% of the donors used for transplant had a positive Hepatitis B surface antigen while the other two cohorts showed 0% positivity (Table 4). However, 3% of the donors belonging to the research cohort and 1% of the donors of the transplant cohort showed positive Hepatitis B Core antigen antibody, indicating a previous Hepatitis B infection.

**Table 4.** Trends in donor serology.

| **Characteristics** | **Research Lungs**<br>**(n = 128)** | **Transplanted lungs**<br>**(n = 391)** | **Rejected lungs**<br>**(n=1167)** |
| --- | --- | --- | --- |
| **Hepatitis B** |  |  |  |
| *Nucleus Amplification Test* |  |  |  |
| Reactive | 0% | 0% | 0.26% |
| Non-Reactive | 100% | 100% | 99.74% |
| *Anti-Hepatitis B core antigen antibody* |  |  |  |
| Reactive | 3.23% | 1.02% | 4.20% |
| Non-Reactive | 96.87% | 98.98% | 95.80% |
| *Hepatitis B surface antigen* |  |  |  |
| Reactive | 0% | 0.26% | 0.43% |
| Non-Reactive | 100% | 99.74% | 99.57% |
| **Hepatitis C** |  |  |  |
| *Nucleus Amplification Test* |  |  |  |
| Reactive | 0% | 4.09% | 7.97% |
| Non-Reactive | 100% | 95.91% | 92.03% |
| *Anti-Hepatitis C antigen antibody* |  |  |  |
| Reactive | 0% | 7.67% | 12.17% |
| Non-Reactive | 100% | 92.33% | 87.83% |
| **HIV\*** |  |  |  |
| *Nucleus Amplification Test* |  |  |  |
| Reactive | 0% | 0% | 0.26% |
| Non-Reactive | 100% | 100% | 99.74% |
| *Anti-HIV antigen antibody* |  |  |  |
| Reactive | 0% | 0% | 0.43% |
| Non-Reactive | 100% | 100% | 99.57% |
| **CMV\*** |  |  |  |
| Reactive | 54.68% | 55.75% | 54.58% |
| Non-Reactive | 45.31% | 44.25% | 45.42% |

\*Abbreviations: HIV – Human Immunodeficiency Virus; CMV – Cytomegalovirus.

8% of the donors whose lungs were rejected and 4% of the donors whose lungs were transplanted showed a reactive Hepatitis C Nucleic Acid Amplification Test (NAAT). None of the lungs used for research had NAAT positivity in the donor, nor a positive Hepatitis C antigen antibody (Table 4). None of the donors belonging to both research and transplant cohorts showed reactivity to HIV NAAT test and HIV antigen antibody testing.

### Microbiological culture data

The most prevalent micro-organism detected on respiratory cultures among all three cohorts was Methicillin Sensitive Staphylococcus Aureus (MSSA) approximately comprising 23% in lungs used for research and transplantation (Table 5). Among Gram negative organisms isolated from lungs used for research, Hemophilus influenzae was seen in 6%, Enterobacter and Klebsiella were observed 4%.

**Table 5.** Trends in microbial respiratory and blood testing.

|  | **Research Lungs**<br>**(n = 128)** | **Transplanted lungs**<br>**(n = 391)** | **Rejected lungs**<br>**(n=1167)** |
| --- | --- | --- | --- |
| **RESPIRATORY TESTING** |  |  |  |
| **Gram positive Bacteria** |  |  |  |
| *MRSA* | 6.25% | 6.14% | 7.20% |
| *MSSA* | 22.66% | 23.27% | 19.40% |
| *Other sp.* | 0% | 0% | 0.77% |
| *Other Staphylococcus sp.* | 0.78% | 0% | 0.42% |
| *Other Streptococcus sp.* | 4.69% | 8.18% | 3.94% |
| *Streptococcus pneumoniae* | 3.13% | 4.35% | 3.85% |
| **Gram Negative Bacteria** | 0% |  |  |
| *Acinetobacter sp.* | 0% | 1.02% | 1.71% |
| *Escherichia coli* | 1.56% | 1.02% | 2.14% |
| *Enterobacter sp.* | 3.91% | 6.14% | 3.08% |
| *Hemophilus influenzae* | 6.25% | 6.91% | 6.16% |
| *Klebsiella sp.* | 3.91% | 2.81% | 3.68% |
| *Other sp.* | 7.03% | 4.60% | 4.37% |
| *Pseudomonas sp.* | 2.34% | 1.02% | 3.59% |
| **Viruses** |  |  |  |
| *SARS COV-2* | 0% | 0.26% | 1.02% |
| *Rhinovirus* | 0.78% | 0.77% | 1.28% |
| *Influenza* | 1.56% | 0% | 0.34% |
| *Parainfluenza* | 0% | 0% | 0% |
| *Adenovirus* | 0% | 0.26% | 0.25% |
| *Enterovirus* | 0.78% | 0.77% | 1.03% |
| *Other virus* | 1.56% | 0% | 0.09% |
| **Fungi** |  |  |  |
| *Aspergillus* | 0.78% | 0.26% | 0.09% |
| *Candida* | 0.78% | 0.51% | 0.69% |
| *Yeast* | 0% | 1.28% | 1.37% |
| *Other fungi* | 0% | 0% | 0.43% |
| **BLOOD CULTURES** |  |  |  |
| **Bacteremia** |  |  |  |
| *MSSA* | 3.9% | 2.30% | 4.37% |
| *MRSA* | 0.79% | 0% | 1.37% |
| *Streptococcus sp.* | 2.34% | 2.30% | 1.88% |
| *Staphylococcus sp. other than MSSA.* | 3.13% | 7.16% | 7.54% |
| *Other Gram-positive sp.* | 0.78% | 0.51% | 0.94% |
| *Pseudomonas* | 0% | 0% | 0.17% |
| *Klebsiella* | 0% | 0% | 0.17% |
| *Other Gram-negative sp.* | 0.78% | 0.77% | 0.90% |
| **Fungemia** |  |  |  |
| *Candida sp.* | 0% | 0% | 0.09% |
| *Other fungi sp.* | 0% | 0% | 0% |

\*Abbreviations – MRSA – Methicillin resistant *Staphylococcus aureus*; MSSA – Methicillin sensitive *Staphylococcus aureus*; sp. – species.

Influenza was the most common virus detected in respiratory cultures of lungs used for research (1.5%). The proportion of Pseudomonas in respiratory cultures was highest among the lungs that were rejected (4%). The proportions of rhinovirus, adenovirus and fungi were <1%.

### Radiographical findings

Clear chest radiographs (scored 0) were only observed in 16% of donors used for research, as compared to donors whose lungs were transplanted (40%). Chest radiographs having opacities in > 1 lobe (scored 3) were highest in donors whose lungs were rejected at 38%, closely followed by 36% in donors whose lungs were utilized for research and only 9% of the donors whose lungs were transplanted (Table 6).

**Table 6.**
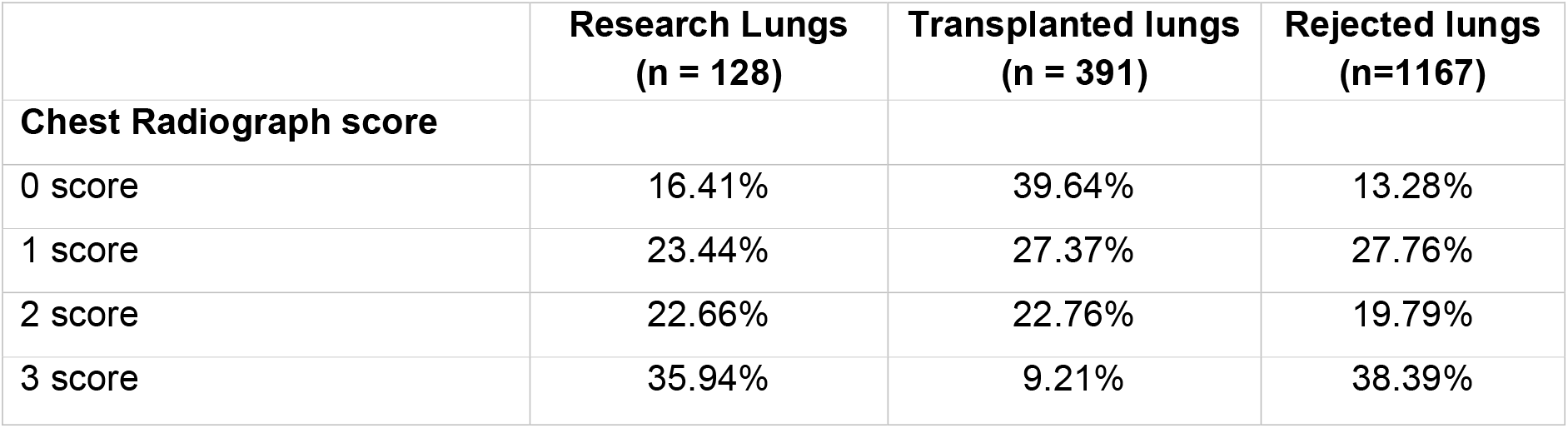
Grading of chest radiographs.

|  | **Research Lungs**<br>**(n = 128)** | **Transplanted lungs**<br>**(n = 391)** | **Rejected lungs**<br>**(n=1167)** |
| --- | --- | --- | --- |
| **Chest Radiograph score** |  |  |  |
| 0 score | 16.41% | 39.64% | 13.28% |
| 1 score | 23.44% | 27.37% | 27.76% |
| 2 score | 22.66% | 22.76% | 19.79% |
| 3 score | 35.94% | 9.21% | 38.39% |

## DISCUSSION

There are strict selection criteria and protocols combined with individualization of decisions which determine acceptance of donor lungs for transplantation. [(9)] However, no such strict selection criteria or protocols exist when human donor lungs are obtained for ex-vivo research. Various ex-vivo studies have advanced our understanding of lung pathophysiology, including understanding immune responses during infection, injury, and repair. [(10),(11)]

A study published in 2020 by the International Thoracic Organ Transplant Registry of the International Society for Heart and Lung Transplantation (ISHLT) mentions that donor variables impacting 1-year and 5-year survival of lung transplant recipients are donor age, history of substance abuse and smoking, diabetes and hypertension across all geographical regions including Europe and North America. [(12)] Similarly, there are various pathophysiological mechanisms by which these above-mentioned variables affect human lung tissue, which can influence the outcomes of ex-vivo lung tissue experiments. [(3),(4)]

Decline in adaptive and innate immunity functions have been noted in lungs obtained from older donors due to factors such as mitochondrial dysfunction, reduced T cell subpopulations, and impaired B cell generation. [(13)] Age also influences bronchoalveolar total cell concentration, neutrophils, and immunoglobin concentration in lung tissue. The CD4+/CD8+ ratio is also significantly increased in the older age group, with an increased ability of alveolar macrophages to release of superoxide in response to a stimuli, resulting in low-grade inflammation in the lungs of donors aged > 65 years old. [(14)] Thus, age of the donor lung tissues should ideally be reported, such as that it can be accounted for as a variable in ex vivo studies.

Diabetes mellitus is known to induce inflammatory as well as fibrotic changes in the lung, which is evidenced by increased levels of inflammatory cytokines like MIP-1δ, IP-10, RANTES, TGF-β1, TNF-α and MIP-1β in lungs of diabetics compared to those of non-diabetics.[(15)] It also leads to a decrease in innate immune responses in the lung, as evidenced by a marked reduction in Toll-like receptor (TLR) protein i.e., of TLR2 and TLR4, along with thickening of alveolar epithelial and capillary basal laminae.[(6),(16)] There is evidence that long-standing diabetes is responsible for the activation of a hyperglycemia-induced pathway resulting in an overproduction of superoxide in the mitochondria by its electron transport chain.[(17)] Thus, more investigation is needed to test the effects of hyperglycemia on pulmonary immune responses.

We noted that at least a quarter of the donors whose lungs had been used for research had a history of smoking. The exposure to cigarette smoke has several effects on the lung. For example, exposure to cigarette smoke accelerates senescence in lung epithelial cells and results in an altered regulation of miRNA, which is an important epigenetic mechanism in aging.[(18)] Due to the impact of smoking on the lung, there is disruption of innate immune responses and induction of a pro-inflammatory state.

Studies have demonstrated altered macrophage phenotype in lungs exposed to cigarette smoke causing inflammation. For example, there is a higher proportion of macrophages with an M2-like polarized phenotype, which is associated with tissue remodeling, upon exposure to cigarette smoke. [(19)] In some individuals exposed to cigarette smoke, the ability of alveolar macrophages to kill bacteria or viruses, and their capacity to remove dead cells is significantly impaired along with chemical modifications of the extracellular matrix (ECM).[(20)]

Further, reduced alveolar macrophage expression is also seen due to decreased expression of macrophage maturation markers like CD44, CD71, CD31 and CD91.[(21)] There is also a loss of mucosal defense by reduction in B cell activating factor belonging to the tumor necrosis factor family thereby reducing levels of secretory IgA. [(22),(23)] Additionally, human bronchial epithelial cells exposed to cigarette smoke show a reduction in IL-8 levels in the supernatant, which is a key mediator in neutrophil chemotaxis when induced by endotoxin or LPS. [(24)] Components of innate antiviral defense mechanisms such as interferons and functions of fibroblasts are also markedly reduced in lungs exposed to cigarette smoke, due to the inability of the human lung fibroblasts to respond to interferon-beta stimulation and epithelial cells to mount an antiviral response. [(25)] Specifically, these cell types show a decrease in expression of interferon-stimulated gene 15 and interferon regulatory factor-7 along with suppression of nuclear translocation of important transcription factors such as nuclear factor-kappa B and interferon regulatory factor-3. [(25)] Thus, a history of current cigarette smoking needs to be considered in ex vivo studies involving human lung tissue.

Our study has certain limitations. Data regarding high-risk behavior and exposure to cigarette smoke (cigarette pack years), cocaine, IV drugs, and non-IV drugs were based on questionnaires filled by either donors or family/friends of donors hence can be subject to recall bias. A dose-response relationship in cigarette smoke is well established and the same applies to various effects on immune responses.[(26),(27)] There is lack of data regarding precise amount of exposure to cigarette smoke measured by pack-years. Similarly, adequacy of blood glucose control in diabetics has differential effect on lung tissue, alveolar and bronchial cells, compared to poorly controlled diabetes.[(28),(29)] We did not have access to glycosylated hemoglobin levels in this study, to account for diabetes control, nor did we have the number of years the donors had diabetes. We also used a simplified radiology score to assess chest radiographs. Computed tomography (CT) imaging and magnetic resonance imaging (MRI) of the lung is superior in assessing quality of lungs prior to donation compared to chest radiographs.[(30)] However, a lack of a standardized tool or a grading method to group CT or MRI findings of lungs donors led to the exclusion of this data which could have potentially provided a better idea on the overall quality of lungs used for research, transplant and those that were rejected.

Considering that lungs from older donors, those with diabetes mellitus or with a history of smoking have significant variations in cellular pathology, genetic composition of alveolar cells, grade of inflammation and varied responses of immunological functions, it becomes crucial to select/match lungs for research based on donor factors, or clearly report these characteristics to the extent of what are available. Moreover, additional research is required to better understand the impact of these donor co-morbidities and exposures on ex-vivo studies of donor lung tissue.

---

**Figure 1:**
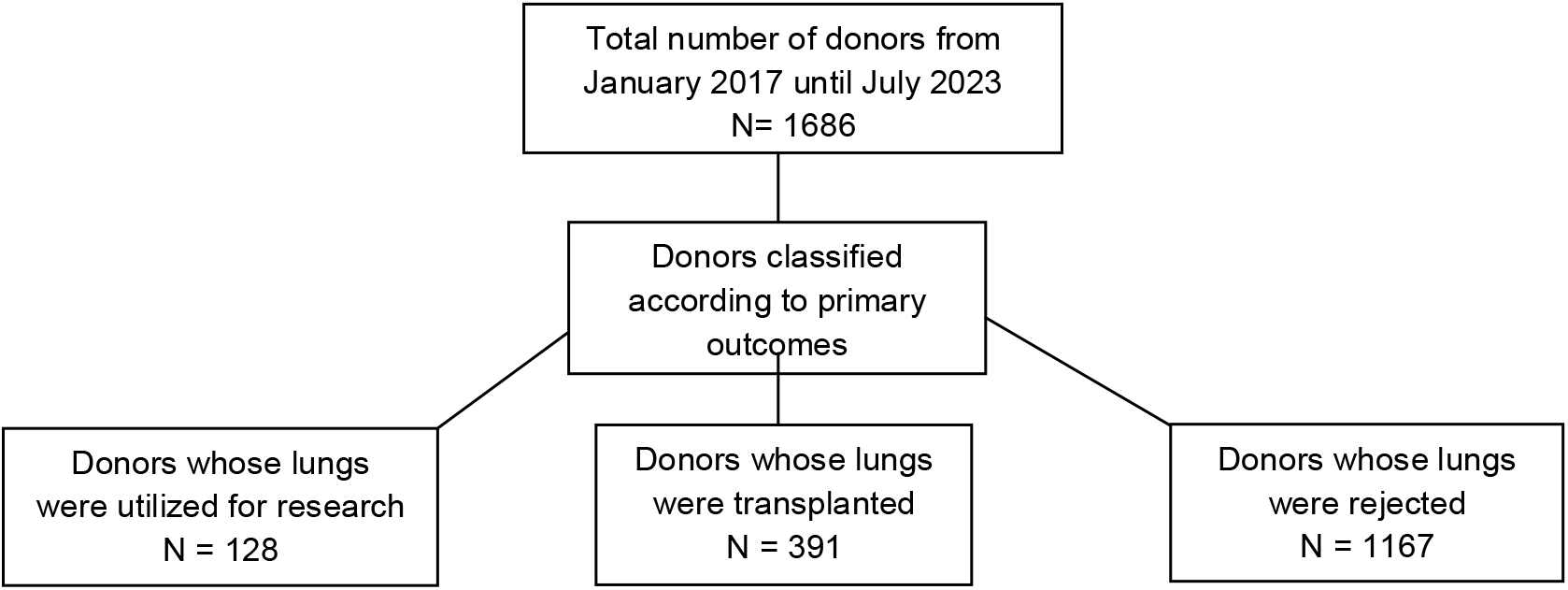
Consort diagram representing primary lung donor characteristics for each primary outcome. **Legend:** We procured data of all lung transplant donors with various outcomes at Mid-America Transplant from January 2017 until July 2023. The lungs were characterized based on their predisposition into three groups - lungs used for research, lungs used for transplant, and lungs which were not recovered from donors.

